# Intranasal oxytocin enhances intrinsic corticostriatal functional connectivity in women

**DOI:** 10.1101/068585

**Authors:** Richard A. I. Bethlehem, Michael V. Lombardo, Meng-Chuan Lai, Bonnie Auyeung, Sarah K. Crockford, Julia Deakin, Sentil Soubramanian, Akeem Sule, Prantik Kundu, Valerie Voon, Simon Baron-Cohen

**Affiliations:** Autism Research Centre, Department of Psychiatry, University of Cambridge, Cambridge, United Kingdom; Center for Applied Neuroscience, Department of Psychology, University of Cyprus, Nicosia, Cyprus; Child and Youth Mental Health Collaborative at the Centre for Addiction and Mental Health and The Hospital for Sick Children, Department of Psychiatry, University of Toronto, Toronto, Canada; Department of Psychiatry, National Taiwan University Hospital and College of Medicine, Taipei, Taiwan; Department of Psychology, School of Philosophy, Psychology and Language Sciences, University of Edinburgh, Edinburgh, United Kingdom; Department of Psychiatry, University of Cambridge, United Kingdom; Cambridgeshire and Peterborough NHS Foundation Trust (CPFT), Cambridge, UK; Brain Imaging Center, Icahn Institute of Medicine at Mt. Sinai, 1470 Madison Ave., 1st floor, New York, NY 10029, USA; Translational and Molecular Imaging Institute, Icahn Institute of Medicine at Mt. Sinai, 1470 Madison Ave., 1st floor, New York, NY 10029, USA; Behavioural and Clinical Neuroscience Institute, University of Cambridge, Cambridge, United Kingdom; National Institute for Health Research Biomedical Research Council, University of Cambridge, United Kingdom; CLASS Clinic, Cambridgeshire and Peterborough NHS Foundation Trust (CPFT), United Kingdom

## Abstract

Oxytocin may influence various human behaviors and the connectivity across subcortical and cortical networks. Previous oxytocin studies are male-biased and often constrained by task-based inferences. Here we investigate the impact of oxytocin on resting state connectivity between subcortical and cortical networks in women. We collected resting state fMRI data on 26 typically-developing women 40 minutes following intranasal oxytocin administration using a double-blind placebo-controlled crossover design. Independent components analysis (ICA) was applied to examine connectivity between networks. An independent analysis of oxytocin receptor (*OXTR*) gene expression in human subcortical and cortical areas was carried out to determine plausibility of direct oxytocin effects on *OXTR*. In women, *OXTR* was highly expressed in striatal and other subcortical regions, but showed modest expression in cortical areas. Oxytocin increased connectivity between corticostriatal circuitry typically involved in reward, emotion, social-communication, language, and pain processing. This effect was 1.39 standard deviations above the null effect of no difference between oxytocin and placebo. This oxytocin-related effect on corticostriatal connectivity covaried with autistic traits, such that oxytocin-related increase in connectivity was stronger in individuals with higher autistic traits. In sum, oxytocin strengthened corticostriatal connectivity in women, particularly with cortical networks that are involved in social-communicative, motivational, and affective processes. This effect may be important for future work on neurological and psychiatric conditions (e.g., autism), particularly through highlighting how oxytocin may operate differently for subsets of individuals.

## Introduction

Oxytocin is a neuropeptide hormone involved in sexual intercourse, childbirth and parent-infant bonding, affecting reward processing, anxiety and social salience ^1^. Oxytocin is not necessarily a ‘pro-social’ hormone, as effects are highly context- and person-dependent^1,2^. Oxytocin has received substantial interest as a potential treatment for psychiatric conditions such as autism spectrum conditions (ASC; henceforth autism) ^3^, although clinical trials show modest effects ^4–6^. Given the marked heterogeneity in autism ^7^ it is possible that the benefits of oxytocin may vary substantially between individuals. For example, on-average intranasal oxytocin improves eye contact during naturalistic social interaction, but the largest effects occur for individuals who typically make the least amount of eye contact ^8^. Thus, in evaluating oxytocin’s therapeutic potential, we must move towards a more precise understanding of how its effects may vary across individuals.

We have theorized that the widespread effects of oxytocin on complex human social behavior may be due to distributed influence at a neural circuit level ^9^. Although oxytocin acts directly at a local level via the oxytocin receptor (*OXTR*), it can potentially affect widespread circuit-level dynamics via connections to areas that are densely populated with *OXTR*. One way to test the hypothesis that oxytocin affects circuit-level organization in the human brain is through oxytocin-administration studies within the context of in-vivo measurement of intrinsic functional brain organization (i.e. connectome or brain network organization) with resting state fMRI (rsfMRI) data. While there are a number of existing neuroimaging oxytocin-administration studies ^9^, most have relied on task-based fMRI paradigms and largely focus on males. In the oxytocin literature there is a prominent bias towards males, and one that affects much of neuroscience and medical research ^10^. Sex differences in the *OXTR* system are documented ^11–13^, suggesting that findings in males may not generalize to females. Furthermore, task-based fMRI has often shown opposite findings in males and females namely in terms of amygdala activation ^14,15^. Because oxytocin is viewed as a potential pharmacotherapy for conditions like autism, and given that sex may play a large moderating roles in drug effectiveness ^16^, it is essential to begin examining how oxytocin operates in the female brain. In addition, although there is a strong male bias in autism diagnoses ^17^ there is reason to believe that females are strongly underrepresented that may have increased the male-biased understanding of autism ^18^. Given the lack of prior literature on oxytocin’s network level effects on brain connectivity in women we chose to use a robust data-driven (hypothesis-free) approach to assess potential connectivity differences.

The majority of studies investigating how oxytocin affects the human brain use task-based fMRI paradigms. While task-based studies are important for targeting specific psychological processes, examination of oxytocin-related effects may, as a result, be neuroanatomically constrained to specific circuits related to those tasks. Examination of functional connectivity using rsfMRI data allows for task-independent assessment of oxytocin’s effect on intrinsic functional brain organization across the entire connectome. Furthermore, the small number of existing rsfMRI oxytocin-administration studies ^5,13,19,20^ use seed-based analyses that do not allow for hypothesis-free examination across the connectome. Thus, a more unconstrained approach could provide novel insights into oxytocin-related effects on connectome organization, especially when little to no prior hypothesis can be derived from existing literature.

Here we use independent components analysis (ICA) to examine how connectivity between-circuits (i.e. between-component connectivity) ^21,22^ differs across oxytocin and placebo. To facilitate our understanding of oxytocin-effects on connectivity in the human brain we analyzed two publicly available post-mortem human brain gene expression datasets to answer the question of how the oxytocin receptor (*OXTR*) is expressed across a variety of subcortical and cortical areas in the human brain. To date, information on OXTR expression has largely been confined to animal studies and translation from that is problematic ^23^. We predicted that oxytocin would have largest impact on connectivity between the densely *OXTR*-populated striatum and cortical circuits. Furthermore, we predicted that impact of oxytocin on connectivity would vary as a function of variation in autistic traits, with larger effects for individuals with higher levels of autistic traits ^8^.

## Methods

### Participants

All research was conducted in accordance with the Declaration of Helsinki and the study had received ethical approval from the NHS Research Ethics Service (NRES Committee East of England – Cambridge Central; REC reference number 14/EE/0202). This study was exempt from clinical trials status by the UK Medicines and Healthcare Regulatory Agency (MHRA).

In a double-blind randomized placebo-controlled cross-over design, 26 women (age: 23.6±4.6 years, range [21-50]) received an oxytocin nasal spray (24 IU, 40.32 µg, Syntocinon-spray; Novartis, Switzerland, pump-actuated) in one session and placebo (the same solution except for the active oxytocin) in the other session in a counterbalanced order. After instruction by a trained medical doctor the sprays were self-administered 40 minutes prior ^24^ to undergoing resting-state fMRI imaging. Participants confirmed no nasal congestion or obstruction on the day of testing. This timing and dosage are by far the most commonly used in oxytocin administration studies to date ^25^. Sessions were separated by at least one week (to ensure full wash-out from the first administration) when participants were on hormonal contraceptive (19/26). When participants were not on hormonal contraceptive (7/26) both sessions took place in the early follicular phase of the menstrual cycle to ensure similar hormone levels between sessions. Exclusion criteria included pregnancy; smoking; a diagnosis of bipolar, obsessive-compulsive, panic or psychotic disorder; use of any psychoactive medication within one year prior to the study; substance dependence; epilepsy; and being post-menopausal. These criteria were assessed by self-report and participants’ general practitioners were given the full protocol prior to participation and asked to notify the research term if they thought there was any reason for exclusion. More details on the testing procedure and sample are provided in the supplementary information (SI) and supplementary table (S1). Briefly, all subject completed the Wechsler Abbreviated Scale of Intelligence^26^ (mean 115.3±13.19), empathy quotient^27^ (mean 55.6±14.53)) and autism quotient^28^ (mean 14.4±7.32) questionnaires prior to the first scanning session. None of the participants had received a formal diagnosis of autism nor did they give any indication that they may have gone undiagnosed. Though we acknowledge no assessment was done to formally rule this out. They were instructed to refrain from alcohol or caffeine on the day of testing and from food and drink 2 hours prior to testing (except for water).

To understand whether oxytocin or some other placebo-related effect that explains any drug-related differences in connectivity, we utilized an independent dataset of age-matched typical females in order to ascertain what are the normative baseline between-component connectivity effects. Our logic here is that if normative connectivity looks similar to patterns we see during placebo, then we can reasonably infer that oxytocin is the primary reason for the induced change in connectivity and not due to some placebo-related change and no effect of oxytocin. This independent dataset consisted of 50 females whom were slightly older but did not statistically differ in age (mean age 31.6±12.2, Wilcoxon rank-sum test: W = 764.5, p = 0.117) collected on the same scanner and which used a similar multi-echo EPI sequence for data collection (see Morris *et al*, 2016 for full details).

### Image acquisition and pre-processing

MRI scanning was done on a 3T Siemens MAGNETOM Tim Trio MRI scanner at the Wolfson Brain Imaging Centre in Cambridge, UK. For the oxytocin-dataset, a total of 270 resting-state functional volumes (eyes-open, with fixation cross) were acquired with a multi-echo EPI ^30^ sequence with online reconstruction (repetition time (TR), 2300 ms; field-of-view (FOV), 240 mm; 33 oblique slices, alternating slice acquisition, slice thickness 3.8 mm, 11% slice gap; 3 echoes at TE = 12, 29, and 46 ms, GRAPPA acceleration factor 2, BW=2368 Hz/pixel, flip angle, 80°). Anatomical images were acquired using a T1-weighted magnetization prepared rapid gradient echo (MPRAGE) sequence (TR, 2250 ms; TI, 900 ms; TE, 2.98 ms; flip angle, 9°; matrix 256×256×256, FOV 256 mm). For the independent rsfMRI dataset on age-matched females, data was acquired on the same 3T scanner and with a multi-echo EPI sequence that was similar to the oxytocin-dataset (TR, 2470 ms; FOV, 240 mm; 32 oblique slices, alternating slice acquisition, slice thickness 3.75 mm, 10% slice gap; 4 echoes at TE = 12, 28, 44, and 60 ms, GRAPPA acceleration factor 3, BW=1698 Hz/pixel, flip angle, 78°). Multi-echo functional images were pre-processed and denoised using the AFNI integrated multi-echo independent component analysis (ME-ICA, meica.py v3, beta1; http://afni.nimh.nih.gov) pipeline ^31^, details on this procedure are outlines in the supplementary information.

### Gene expression analysis

To better characterize subcortical and cortical brain regions in terms of *OXTR* gene expression, we analyzed RNAseq data in the Allen Institute BrainSpan atlas (http://www.brainspan.org) and the Genotype-Tissue Expression (GTEx) consortium dataset (http://www.gtexportal.org/home/). The BrainSpan atlas covers a number of cortical areas the might provide insights into potential cortical targets of oxytocin expression, whereas the GTEx dataset does not have many regionally-specific areas of the cortex (only BA9 and BA24) and mostly includes more detailed information on several subcortical brain regions. In these analyses we used all postnatal (birth to 79 yrs.) samples in each dataset, stratified by biological sex. *OXTR* was isolated and plots were produced to descriptively indicate expression levels across brain regions. Expression levels in both datasets were summarized as Reads Per Kilobase of transcript per Million mapped reads (RPKM). Full details for the BrainSpan and GTEx procedures are available in their white papers; http://bit.ly/2dqRF47 and http://bit.ly/2e8o1W2 respectively. To examine whether *OXTR* expression levels were significantly elevated in each brain region, we compared expression levels against zero and, as a more conservative test, against another tissue from GTEx where we would not expect *OXTR* to be expressed (i.e. skin). These tests were carried out using permutation t-tests (1000 permutations) implemented with the perm.t.test function in R.

### Group Independent Components Analysis and Dual Regression

To assess large-scale intrinsic functional organization of the brain we first utilized the unsupervised data-driven method of independent component analysis (ICA) to conduct a group-ICA followed by a dual regression to back-project spatial maps and individual time series for each component and subject. Both group-ICA and dual regression were implemented with FSL’s MELODIC and Dual Regression tools (www.fmrib.ox.ac.uk/fsl). For group-ICA, we constrained the dimensionality estimate to 30, as in most cases with low-dimensional ICA, the number of meaningful components can be anywhere from 10-30 ^21^. Some components were localized primarily to white matter and although likely may be driven by true BOLD-related signal (due to high ME-ICA kappa weighting), were not considered in any further analyses. 22 out of 30 components were manually classified as primarily localized to gray matter and were clearly not noise-driven components. Correlation matrices were constructed for all component pairs, these were assessed for significance using paired sampled t-tests and resulting p-values were corrected for multiple comparison using Bonferroni correction at a family-wise error rate of 5%. Difference scores were computed for pairs that survived FWE correction on the Fisher z-transformed correlation scores ^32^ and entered into robust regression (for insensitivity to outliers) ^33^ with AQ scores. For more details see the supplementary information.

### Large-Scale Reverse Inference with Cognitive Decoding in NeuroSynth

To better characterize the components showing an oxytocin-related effect on connectivity we used the decoder function in NeuroSynth ^34^ to compare the whole-brain component maps with large-scale automated meta-analysis maps within NeuroSynth. The top 100 terms (excluding terms for brain regions) ranked by the correlation strength between the component map and the meta-analytic map were visualized as a word cloud using the wordcloud library in R, with the size of the font scaled by correlation strength.

## Results

### Oxytocin Receptor (*OXTR*) Gene Expression in the Female Human Brain

Expression profiles of *OXTR* in women derived from the GTEx dataset reveal broad expression across subcortical regions, but with notable enrichments particularly in nucleus accumbens, substantia nigra, and the hypothalamus (Figure 1). All regions showed *OXTR* expression that was significantly above 0 and critically, was also significantly stronger than expression in a tissue we would expect to show little expression (i.e. skin) (SI Table S2). Cortical regions from the BrainSpan dataset also exhibit significant *OXTR* expression (above 0 and when compared to skin; SI Table S2), albeit at much more modest levels than some subcortical regions. This modest degree of *OXTR* expression may be particularly relevant given studies that show broad oxytocin-related effects on complex human social behavior, social communication, and social cognition that affects distributed cortical regions (e.g., superior temporal gyrus, medial prefrontal cortex (MFC)). However, there is a lack of specificity apparent in *OXTR* expression in cortex, as most regions show similar levels of expression. As a whole, these data indicate that oxytocin could have potent direct effects on *OXTR* within subcortical circuitry, particular areas of the striatum and midbrain, but may also have similar *OXTR*-driven effects to a lesser extent across most cortical areas where *OXTR* expression is modest. Given the lack of specificity within cortex, these data also support an approach for examining oxytocin-related effects on intrinsic functional connectivity that examines all between-networks connections, as all may be susceptible to plausible effects. However, given the enrichment particularly in striatal and midbrain regions, it is likely that oxytocin-related effects on connectivity may particularly affect connections between cortex and the densely OXTR-populated striatum and midbrain. We also carried out exploratory analyses on gender differences in OXTR expression and these are included in the supplementary information (Figure S1).

**Figure 1:**
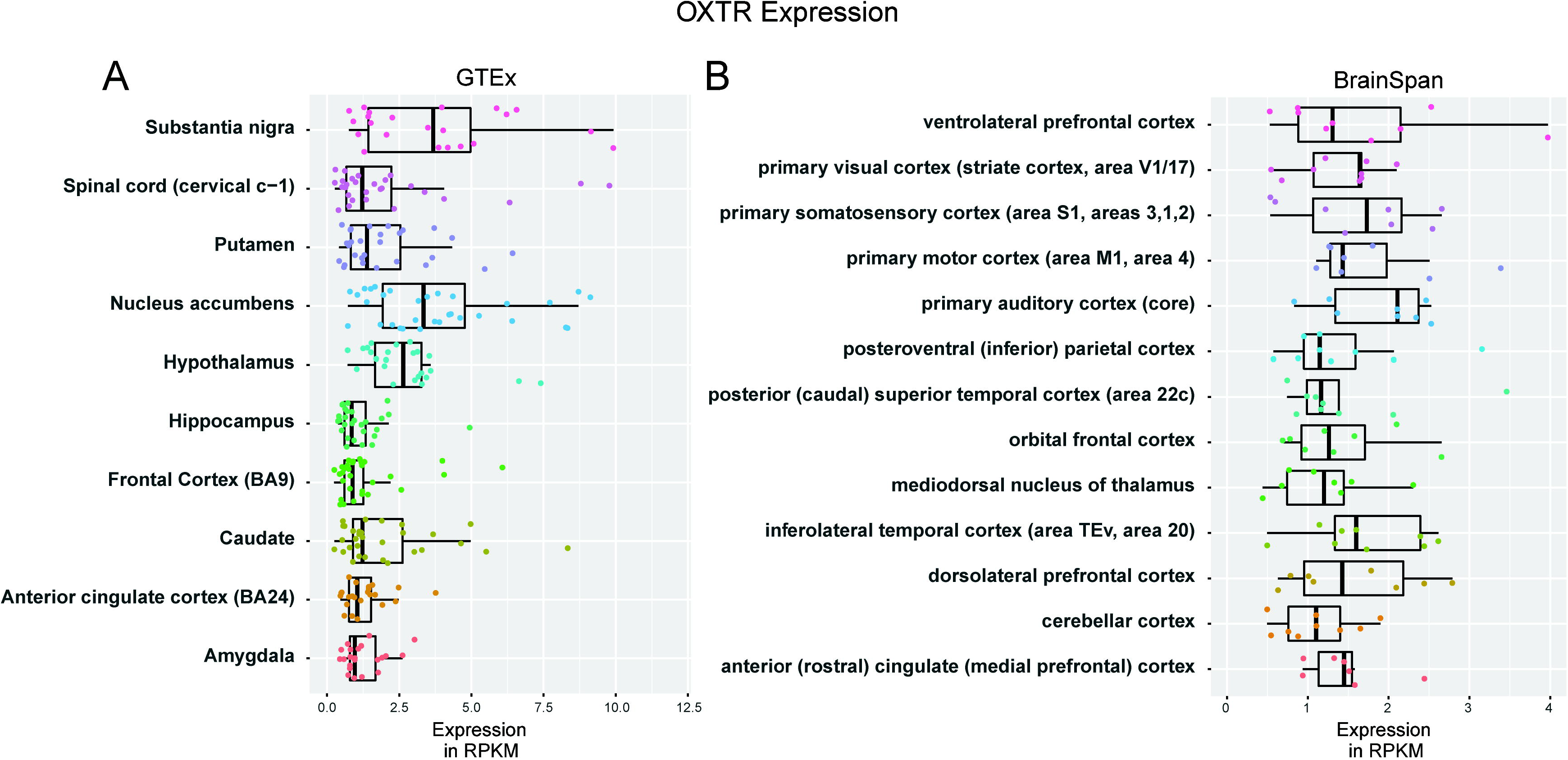
Oxytocin receptor (*OXTR*) gene expression in the female human brain. This figure illustrates *OXTR* gene expression measured via RNAseq in BrainSpan (http://www.brainspan.org) and GTEx (http://www.gtexportal.org/home/) datasets. Panel A shows expression for all subcortical regions available in the GTEx dataset in women. All brain regions show significant expression of *OXTR* above 0 and compared to expression in non-brain (skin) tissue. On-average, *OXTR* expression is particularly enriched in ventral striatum (Nucleus Accumbens), substantia nigra, and hypothalamus. Panel B shows expression for all cortical areas and the thalamus available in the BrainSpan atlas in women. All areas also show significant, albeit modest, levels of *OXTR* expression compared to 0 and non-brain (skin) tissue.

### Oxytocin-Related Between-Component Connectivity Differences

Analyses of all pairwise comparisons of between-component connectivity differences as a function of oxytocin administration revealed only one pair of components, IC11 and IC21 (Figure 2, panels A and B), whose connectivity was substantially affected by oxytocin (*t(24)* = 6.99, *p* = 3.10e-7, effect size d = 1.39, 95% CI [0.96 to 1.86]) and survived after Bonferroni correction (FWE p<0.05) for multiple comparisons. The full pairwise correlation matrix is provided in supplementary figure 2. As shown in Fig 2E, all but 2 participants (92%; 23/25) showed evidence of a non-zero oxytocin-related boost in connectivity over the placebo condition (Figure 2, panel E). Within the placebo condition alone, connectivity was not different from 0 (*t(24)* = –0.86, *p* = 0.39). However, within the oxytocin condition, connectivity was substantially elevated above 0 (*t(24)* = 6.22, *p* = 1.95e-6).

The IC11 component comprised regions in primary auditory cortex, middle and posterior divisions of the insula, superior temporal gyrus, posterior superior temporal sulcus, middle and posterior cingulate cortex, ventromedial prefrontal cortex, amygdala, and superior parietal lobe. These brain regions overlap with areas typically considered important in processes such as language, social-communication, self-referential and social cognition, pain, and emotion ^35–39^. NeuroSynth decoding revealed that most of the terms with the highest correlation with IC11 were predominantly terms referring to pain-related, motor-related, or language/speech-related processes (Figure 2, panel C). The IC21 component was comprised entirely of subcortical regions such as the striatum, basal ganglia, amygdala, thalamus, midbrain, and brainstem. These regions, particularly the striatum, midbrain, and amygdala, are typically considered highly involved in reward and emotion-related processes ^40–42^. This was again confirmed with NeuroSynth decoding showing a predominance of reward, motivation, and affective terms (Figure 2, panel D).

Next, we examined whether individual differences in autistic traits accounted for variability in oxytocin-related effects on connectivity between these networks in an exploratory analysis. Given prior work suggesting that oxytocin may have its largest effect on individuals who show the most atypical social behavior ^8^, we hypothesized that oxytocin may have the largest effects on connectivity in individuals with the highest degree of autistic traits. Here we found evidence confirming this hypothesis, as oxytocin’s effect on between-component connectivity appeared to increase with increased degree of autistic traits: r = 0.41, one-tailed *p = 0.0351* (Figure 2, panel F).

**Figure 2:**
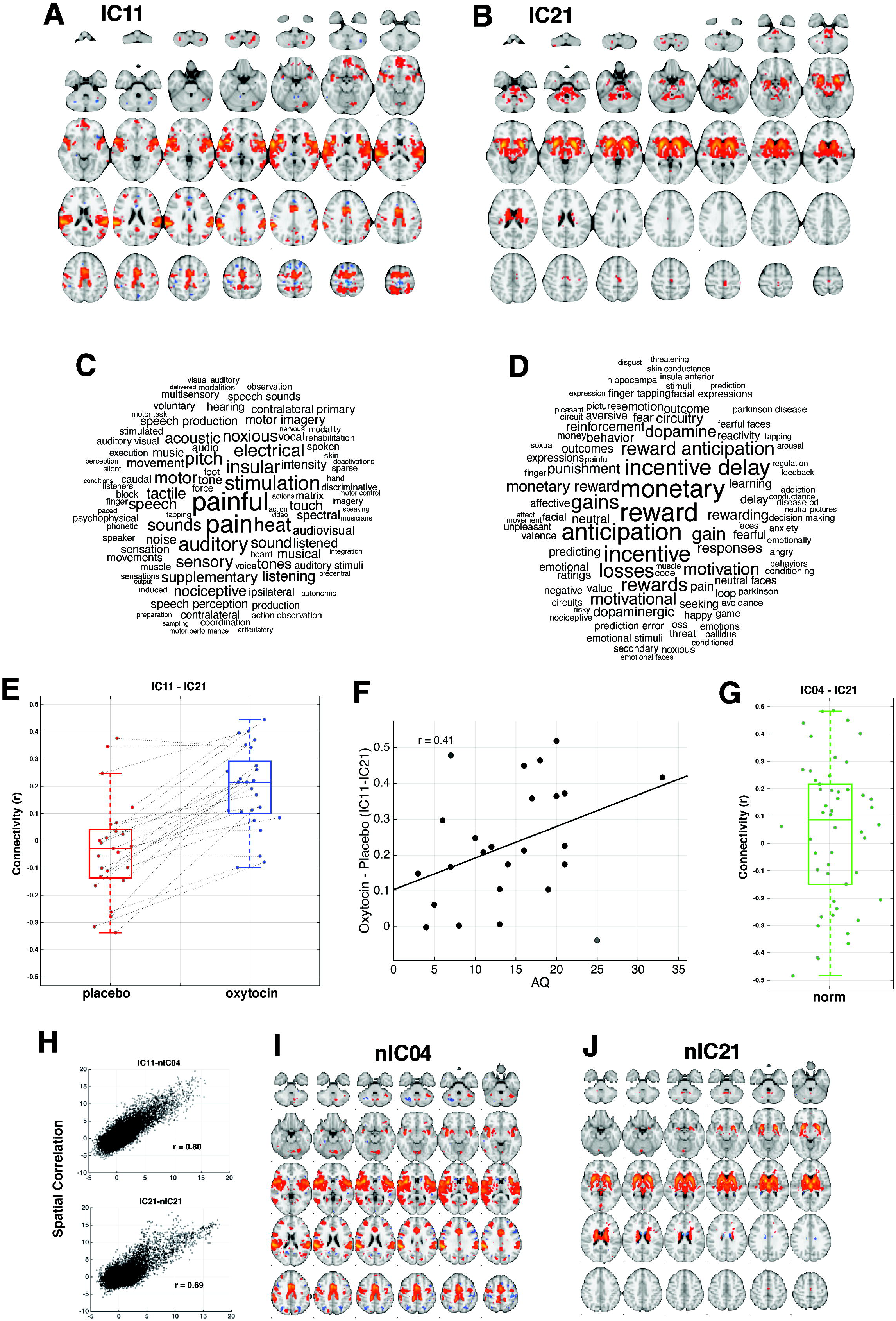
Oxytocin-related enhancement of intrinsic functional connectivity. **Panel A** shows the spatial map of component IC11. Voxels are colored by Z-statistics indicating how well each voxel’s time series fits the component’s time series. **Panel B** shows the same information for component IC21. **Panel C** shows the top 100 terms associated with component IC11 based on NeuroSynth decoding and font size represents relative correlation strength of that term to the component. **Panel D** shows the same information for component IC21. **Panel E** shows connectivity between IC11 and IC21 for each subject during placebo or oxytocin administration. Dots represent individual subjects and the lines connect each individual’s data under placebo and oxytocin, with the positive slopes indicating an enhancement of connectivity after oxytocin administration. Underneath the individual-level data are boxplots that indicate the median, interquartile range, and outer fences. Interestingly, the two individuals who would be considered outliers in the placebo condition are the minority of individuals showing no enhancement of connectivity as a function of oxytocin. **Panel F** shows the relationship between oxytocin-related effects on connectivity and continuous variation in autistic traits as measured by the AQ. **Panel G** shows the between component connectivity of between comparable components of the normative dataset. **Panel H** shows the spatial correlation between the oxytocin data components and the two normative components that were selected. **Panels I and J** show the normative components spatial maps.

Finally, we ran further analyses to aid the interpretation of such an effect. One interpretation could be that oxytocin is the primary driver of enhanced connectivity between these components. However, the alternative could be that oxytocin has no effect on connectivity, and that the placebo might somehow induce a decrease in connectivity between these components. To tease apart these different interpretations, we looked to an independent dataset of rsfMRI to ascertain what the normative connectivity strength is between these two components. If oxytocin was truly enhancing connectivity between these components, we would predict that connectivity between these components under normative conditions would be similar to those seen under placebo. That is, normative connectivity effects between these components should manifest similarly to placebo and on average show no difference from 0. We identified two components that spatially appeared nearly identical to the component pair we observed an oxytocin effect in; nIC4 and nIC21 (Figure 2I & 2J). Quantitatively confirming this similarity, we find very large correlations between the spatial component maps of the normative and oxytocin/placebo datasets (nIC4-IC11, r = 0.80; nIC21-IC21, r = 0.69, see Figure 2H). No other components showed anywhere near such strong correlations (all r < 0.2). Similar to our placebo condition, this component-pair showed connectivity that was not significantly different from zero: t(49) = 1.23, p = 0.22 (Figure 2G). Furthermore, comparison between normative connectivity and connectivity during placebo revealed no statistical difference (t-test with unequal variance assumed and degrees of freedom estimated using Satterthwaite’s approximation; t(64.6) = −1.507, p = 0.1370). This further clarifies our interpretation that it is indeed the oxytocin condition that drives enhancements in connectivity between these components and that the placebo condition is a good approximation of normative functional connectivity effects within this corticostriatal circuit.

## Discussion

This is the first study to investigate how oxytocin affects intrinsic functional organization of the human brain at the level of between-network interactions. We discovered a specific corticostriatal network implicated in social-communicative, motivational, and affective processes that is heavily affected by oxytocin. Under oxytocin the connectivity between these two components was substantially elevated on-average and an oxytocin-related boost was observed in almost all participants. The fact that these corticostriatal connections are not particularly strong under normative conditions and with the administration of placebo, but become increasingly coordinated under oxytocin may be important for understanding how oxytocin influences cognition and behavior. Future work is needed to examine oxytocin-related strengthening of connectivity between these circuits and its effect on specific cognitive and behavioral processes. For example, these corticostriatal connections under pain or social-communication processes may illuminate important brain-behavior links that are affected by oxytocin. These results also illustrate how oxytocin is likely to extend beyond certain brain regions traditionally thought to be important ^43^. For example, previous neuroimaging studies in humans have largely focused on amygdala-related effects and to a lesser extent on striatal regions. The current study suggests oxytocin’s effects may extend well beyond the amygdala and striatum, and most importantly, may incorporate interactions between subcortical striatal regions with cortical areas.

The degree to which oxytocin enhanced connectivity was also associated with continuous variation in autistic traits, such that those with the highest levels of autistic traits showed the largest oxytocin-related effect on connectivity. These results may point towards the idea that oxytocin may have varying impact on different subsets of individuals. Individuals with the highest levels of autistic traits seem to show the largest oxytocin-related connectivity boost. It will be important to extend these ideas into neuropsychiatric conditions such as autism spectrum conditions (ASC). Oxytocin is hypothesized to be of some potential value therapeutically for autism ^43^. However, given the large degree of heterogeneity in ASC ^7^ and the knowledge that therapies may work well for some individuals and not others, it will be of the utmost importance to examine how oxytocin may or may not work well on specific subsets of affected individuals.

Supporting the plausibility of oxytocin-related effects on connectivity between these circuits, we also showed evidence supporting the idea that many of the brain regions involved in both IC11 and IC21 maps show some degree of *OXTR* expression. For instance, it is well known from non-human animal work that the striatum and regions within the midbrain are highly populated with oxytocin receptors ^44,45^. Here we confirmed such finding with evidence from *OXTR* expression in the brain in human females and furthered a proof-of-concept evidence that oxytocin may leverage this enrichment in *OXTR* to influence neural circuits connected to the striatum. We also discovered that there are modest levels of *OXTR* expression throughout many cortical areas. Given the lack of cortical specificity for *OXTR* enrichment, it remains possible that the observed connectivity effects with rsfMRI may not necessarily be mediated by direct action of oxytocin on *OXTR* in specific cortical regions. Rather, oxytocin could exert such effects via other indirect routes, perhaps originating in striatal circuitry where there is highly enrichment in *OXTR* or via other mechanisms of action ^9^. Although expression patterns of *OXTR* were not specific to cortical regions it may be that more fine-grained spatial maps of *OXTR* might provide a clearer picture. For example, the development of a PET ligand could certainly further advance our understanding of *OXTR* distribution in-vivo in the human brain.

This study has several novel elements that need to be highlighted. Specifically, this study focusses specifically on oxytocin-related resting-state effects in women. There are notable male biases throughout neuroscience and medical research and this bias may explain why studies looking at the effects of drugs tend to miss many adverse effects or show a lack of efficacy when applied to females ^10,46^. This bias can be observed in much of the prior work on oxytocin in humans as well, with some neuroimaging studies indicating potential differences in the oxytocin system between sexes ^13,14,47–49^ Part of this bias in oxytocin research might be explained by the higher risk of side-effects (e.g. lactation in mothers, abnormal uterine contractions and elevated blood pressure), though the intransal administration has proven to be a safe methods of administration ^25^. There have been a few studies that assessed the effect of gender in oxytocin administration. For example, previous studies examining functional connectivity during tasks show enhanced connectivity in women but decreased connectivity in men ^47,50^. Although our study was not explicitly set to examine sex differences in the effects of oxytocin, future research should focus on how oxytocin may have different effects across males and females.

Second, surpassing much of the existing neuroimaging work on oxytocin, our study is the first to take a whole-brain, unsupervised approach to examine connectivity between neural networks. The small number of studies examining in-vivo oxytocin-related changes to functional connectivity in humans utilized a seed-based connectivity approach. This approach elucidates effects of oxytocin on connectivity with the pre-selected seed region, but is limited by the a priori selection. As we have shown with the analysis of OXTR expression much of the prior work is not necessarily informed by this expression pattern. It partly lacks specificity for certain regions and the present data suggest that other brain regions, not traditionally reported in oxytocin administration literature, might also have OXTR expression that would make it a potential target for administration. Rather, prior work tends to be heavily directed to regions that are justified based on their role in psychological processes that are linked to oxytocin (e.g., amygdala). In our work, we have taken an unbiased approach to provide insight into oxytocin’s effect on corticostriatal connectivity. These circuits might not have been identified with an approach constrained by task-based activation or seed-based connectivity based on this task-related activation.

The highlighted effect places emphasis on striatal interactions with cortical areas that are associated with pain processing. These results are interesting in light of work showing that oxytocin can not only act as an anxiolytic ^51,52^, but can also act as a painkiller ^53^. It should be noted that the anxiolytic effect might not fully explain oxytocins effect on behavior as benzodiazepines do not show a similar behavioral effect. To our knowledge, there is little neuroimaging work in females focusing on oxytocin and its influence on neural systems for pain processing, as most published work is exclusively on males and/or is focused on empathy for pain ^54–57^. Similar to oxytocin research, research on pain has traditionally been heavily male biased ^16^, while women tend to suffer more from acute and chronic pain ^58^. Our results suggest that future work is needed in this area, particularly on oxytocin’s effect on pain and how such corticostriatal networks may be involved. In addition, areas identified by this data-driven approach show that key areas in the brains reward circuitry are modulated by oxytocin administration. It has been previously hypothesized that oxytocin exerts its effect on social salience and social cognition by modulating stress and reward processing ^1^. The present study also highlights neural systems underlying these cognitive processes as key target for oxytocin administration. Further research into how oxytocin specifically modulates social reward processing might shed further light on its potential to more broadly modulate social cognition.

There are some caveats and limitations to keep in mind. First, the sample size is moderate and potentially provides low power to detect small effects. However, our multi-echo fMRI approach is a strength that could help counteract issues associated with statistical power. Multi-echo EPI acquisition and the ME-ICA denoising technique employed here is known to greatly enhance temporal signal-to-noise ratio (tSNR) and allow for enhanced ability to reduce false positives ^31^. These enhancements tied to principled elimination of non-BOLD noise in rsfMRI could be beneficial for power because reduction in noise potentially increases observable effect sizes ^59^, and reduce effect size estimates for false positive effects. Future work collecting larger samples to replicate and extend these findings would be facilitated by characterizing individuals in continuous variation in autistic traits. Our study indicates that oxytocin-related effects tend to be stronger in individuals with more autistic traits. As noted in the points about sex and gender, future work should also examine whether similar or different effects are present in males. It would also be important to further extend this work in clinically diagnosed individuals with autism. Our exploratory analysis revealed a potential correlation with autistic traits which may suggest that oxytocin could facilitate corticostriatal connectivity in clinically diagnosed patients. If such a relationship extends into the clinically diagnosed population of the autism spectrum, we may expect to see that oxytocin provides the largest enhancements to the most affected individuals ^8^. This should however at this point be considered exploratory.

Furthermore, there has been some debate in recent years about the extend to which oxytocin crosses the blood brain barrier ^60^. A recent study that assessed CSF and plasma concentrations after intranasal administration found elevated levels in plasma after 15 minutes and a peak in CSF elevation at 75 minutes ^61^. By far most studies have used the 24IU dose and timing of 40 minutes to show behavioral effects ^9,25,43^. Yet, is is possible some behavioral effects might originate from peripheral elevation as opposed to a central effect ^60^. Nonetheless, a recent review on the issue suggest that the intranasal route is likely still the best candidate for administration and found no effects from intravenous administration ^62^.

The relation between increased CSF oxytocin and timing of potential behavioral effect also remains unclear. The present study was not set out to determine the best dose or timing or to assess whether oxytocin could cross the blood-brain barrier. Unfortunately, there is currently no PET-ligand available to definitively assess the timing and central binding of intranasal oxytocin, though animal work on this is progressing ^63,64^. Thus, in order to be able to compare the present findings to existing literature, we chose to use the same timing and dosage.

Finally, underpowered studies are common amongst oxytocin administration studies ^65^. The observed effect here between IC11 and IC21 is large. For the current sample size, the minimum effect size achieving 80% power at an alpha of 0.05 is d = 0.6. An effect this low or lower was never observed in our bootstrapping analysis to estimate variability in the IC11-IC21 effect (see Supplemental Figure 3). Therefore, we can reason that we had sufficient power to detect such an effect at the current sample size. As for other more subtle effects, our report here is likely underpowered to detect such effects and much larger studies are likely needed to detect such smaller effects. Since this is the first work on the topic of between-component connectivity in a female rsfMRI oxytocin administration study, we have provided effect sizes estimates for all IC comparisons to aid others in future power calculations (Supplementary Figure 4). Each of the corresponding 22 IC maps can be viewed and downloaded on NeuroVault (http://neurovault.org/collections/2154/).

In conclusion, we have discovered that oxytocin enhances corticostriatal connectivity in women. These corticostriatal networks play roles in social-communicative, motivational, and affective processes and the results may be particularly important for understanding how oxytocin changes neurodynamics that may be relevant for many neuropsychiatric conditions with deficits in those domains and neural circuits. Future work examining these effects in males as well as clinically diagnosed samples will be important, as will be the examination of what subsets of individuals may benefit most from oxytocin-related changes in between-network connectivity.

## Acknowledgements

We would like to thank John Suckling and Pradeep Nathan for valuable discussions during this study. During this research RB was funded by the MRC UK, the Pinsent Darwin Trust and the Cambridge Trust. M-CL is supported by the William Binks Autism Neuroscience Fellowship, Cambridge, and the O’Brien Scholars Program within the Child and Youth Mental Health Collaborative at the Centre for Addiction and Mental Health and The Hospital for Sick Children, Toronto. SB-C is supported by the MRC UK, the Wellcome Trust and the Autism Research Trust. The research was supported by the National Institute for Health Research (NIHR) Collaboration for Leadership in Applied Health Research and Care East of England at Cambridgeshire and Peterborough NHS Foundation Trust. The views expressed are those of the author(s) and not necessarily those of the NHS, the NIHR or the Department of Health.

